# Identifying components that vary in space and time from resting-state functional MRI

**DOI:** 10.1101/2021.07.01.450675

**Authors:** Gregory Scott, Rob Leech

## Abstract

A widespread assumption of fMRI-derived large-scale intrinsic connectivity networks (ICNs) is that they are spatially static over time. However, the assumption of spatial stationarity of ICNs has been challenged by a range of techniques that allow for time-varying connectivity between brain regions and demonstration that canonical networks like the default model network (DMN) can be fractionated according to time-varying connectivity relationships of their subcomponents. Previously, we developed a simple spatiotemporal ICA (stICA) technique to allow the discovery of patterns of spatiotemporal evolution in task fMRI data in a way that avoided the traditional constraint of spatial stationarity on brain networks, and we validated the approach in fMRI of task-to-rest transitions. Here, we apply our stICA technique to resting-state fMRI datasets to explore whether spatiotemporally evolving components of brain activity can be identified in the absence of an overt behavioural task. We found that stICA components could generally be described in terms of graded onsets and offsets of ICNs that had been calculated based on techniques that assumed spatial stationarity. Our results suggest that, to a reasonable approximation, stable ICNs can be taken to be building blocks of the spatiotemporal patterns measured with resting-state fMRI.

## Introduction

In cognitive neuroscience, human brain function is frequently described in terms of the coordinated activity of large-scale functional connectivity networks, or intrinsic connectivity networks (ICNs), that can be identified in functional MRI (fMRI) (Smith et al., 2009). Spatial maps of these ICNs are readily defined using both task and resting-state fMRI data and a range of techniques, historically, seed-based correlational approaches (Biswal et al., 1995) and, more recently, data-driven techniques including spatial independent component analysis (ICA) (McKeown et al., 1998). For many cognitive neuroscientists, ICNs have come to be viewed as the fundamental units of cognitive processes (Bressler and Menon, 2010; Hampshire et al., 2012). A common assumption of this model is that these brain networks are spatially static over time. For example, activity within posterior cingulate, ventromedial prefrontal, and lateral parietal cortices, forming the default mode network (DMN), is considered to be collectively increased at rest, but deactivated during externally-orientated cognitive tasks (Buckner et al., 2008), whereas the frontoparietal control network (FPCN) is said to activate, along with sensory and motor systems, during externally-orientated tasks, and shows a relative deactivation at rest (Dosenbach et al., 2007). However, the assumption of spatial stationarity of ICNs has been challenged by a range of techniques that allow for time-varying connectivity between brain regions (Lurie et al., 2020), and demonstration that canonical networks like the DMN can be fractionated according to differences in the time-varying connectivity relationships of their subcomponents (Leech et al., 2011).

Previously, we developed a data-driven technique to allow the discovery of common patterns of spatiotemporal evolution in fMRI data in a way that avoided the traditional constraint of spatial stationarity on brain networks (Scott et al., 2015). This method combined a simple data reorganization with spatial ICA, enabling a form of spatiotemporal ICA (stICA). When applied to fMRI data from the boundaries of alternating rest and task blocks, our technique identified a spatiotemporal component associated with rest offset-task onset boundaries that evolved from a DMN-like network becoming less active over time to a FPCN-like network becoming more active. We made other observations relevant to understanding the complexities of brain networks, for example, a DMN-like component that evolved spatially, shifting to an anterior/superior pattern of deactivation with task onset from a posterior/inferior pattern during rest. Combined with additional validation using simulated data, these findings suggested our stICA approach is able to identify components that can evolve spatially over time, without having to specify networks *a priori* or assume spatial stationarity of networks.

Here, in an initial application of our stICA approach to resting-state fMRI data, we explore whether spatiotemporally evolving patterns of brain activity consistently arise in the absence of an overt behavioural task. Our approach can detect spatial non-stationarity; however, it does not require it. As such, we can use the technique to examine the extent to which resting-state fMRI is spatially non-stationary, and the extent to which canonical ICNs reflect underlying macroscopic neural dynamics. We focus on the characterisation of the spatiotemporal components identified by our technique, and use the ICN parcellation of Yeo et al (Yeo et al., 2011) to provide a convenient reference framework with which to describe in a quantitative way the spatiotemporal evolution of the resulting components.

## Methods

### Spatiotemporal ICA approach (TARDIS)

We have previously introduced our stICA approach and its application to simulated and empirical task fMRI data (Scott et al., 2015). We herein refer to this method as TARDIS (after Time And Relative Dimension In Space, from the BBC television series Doctor Who, https://bit.ly/30g3AvV). We first summarise the steps of the original TARDIS approach (**Figure 1A**) before describing the simple modification for resting-state fMRI used here (**Figure 1B**): (1) We define anchor points, i.e., theoretically interesting timepoints in a 4D fMRI input dataset (i.e. volume indices); in our previous work these were the times of the onset and offset of task blocks (interleaved with rest blocks). (2) For each input fMRI dataset, we extract the fMRI volume at each anchor timepoint and the following nine timepoints, i.e. a window of 10 fMRI volumes. Note that the length of this temporal window can be modified depending on the neurobiological question and temporal resolution, but we here do not explore other values. (3) We spatially concatenate the 10 volumes extracted for each anchor point along the medial–lateral dimension (although the spatial dimension over which data is concatenated has no effect on the results; we use the medial–lateral dimension for illustrative convenience). This procedure, for example, on 4mm fMRI volumes of image dimensions 23 × 28 × 23 voxels, yields samples each of dimensions 230 × 28 × 23 voxels. (4) The samples (each a 10 volume-wide image) are then concatenated along the fourth dimension (this dimension normally referred to as the “temporal dimension” in fMRI data). This temporal concatenation is done for each subject’s data independently. (5) The resulting 4D datasets are decomposed into independent components by performing a probabilistic ICA in FSL MELODIC (Jenkinson et al., 2012), using the temporally-concatenated group-ICA method. We here limit the analyses to 50 dimensions (components); we have previously shown our task-based findings were robust to different dimensionalities of decomposition. The resulting spatiotemporal ‘TARDIS components’ – as we refer to them here – output by MELODIC are still viewable with conventional tools (e.g. the FSL MELODIC web report, or the `fsleyes` command), each appearing as 10 brains in a row, the medial-lateral dimension signifying time, whereby the left-most brain indicates the first timepoint of the component and the right-most brain the last.

**Figure 1.**
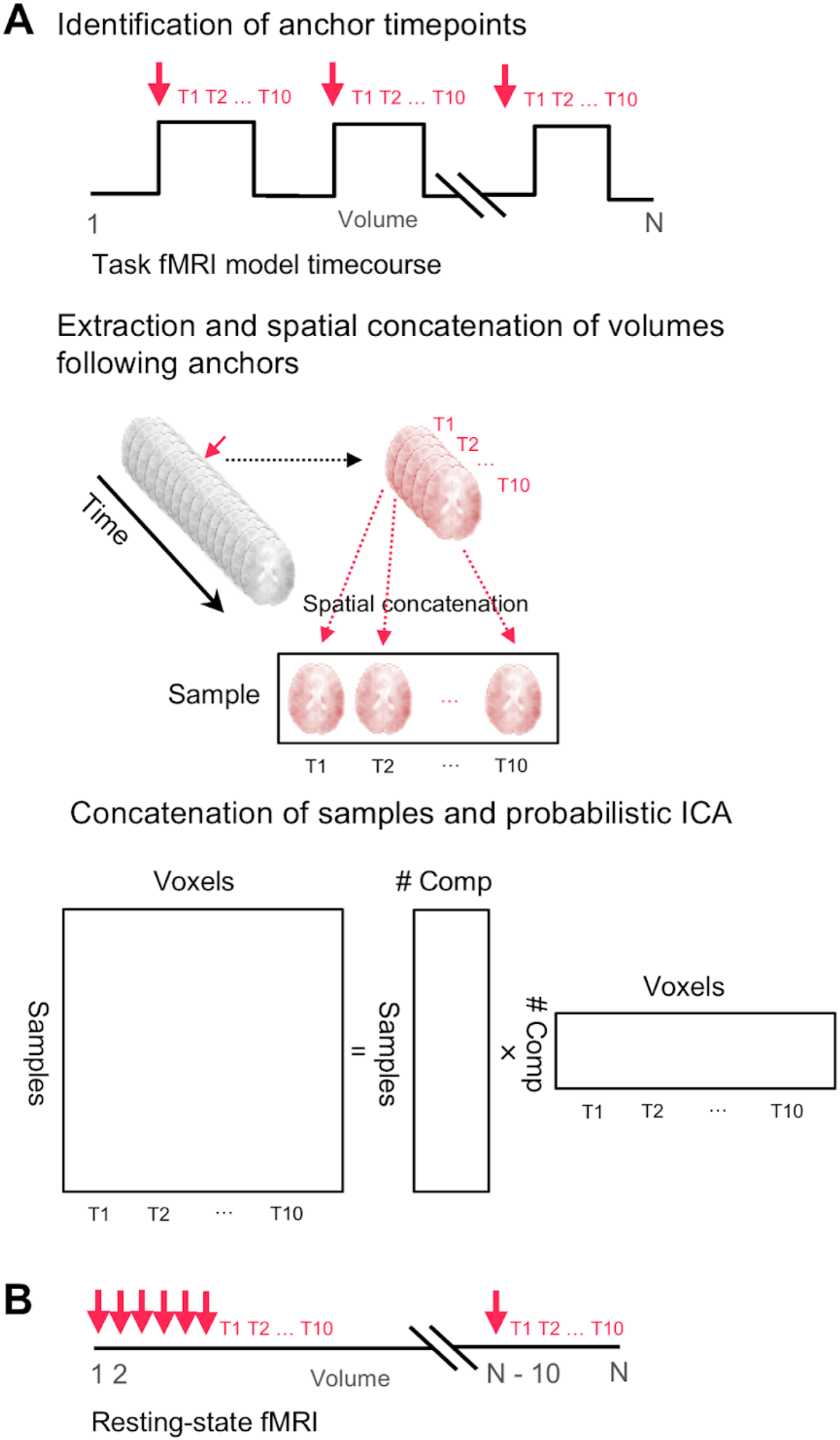
Methodology of the TARDIS approach. **(A)** The original TARDIS approach of spatiotemporal ICA (stICA). Anchor points of theoretical interest are identified, such as marking the timepoint at the start of task blocks. The subsequent N volumes following each anchor point (here, N=10) are then extracted and spatially concatenated along the medial-lateral dimension (although the choice of spatial dimension is arbitrary). The 10-volume wide sample for each anchor point is then concatenated in the fourth dimension (i.e., what is normally thought of as the temporal dimension), and the data for all subjects are then entered into a group ICA (MELODIC). **(B)** The method is modified for resting-state by defining anchor points instead sequentially throughout the fourth dimension of the fMRI data (i.e., first to last-10 timepoints). The steps then proceed as for (A).

Here, we extend the TARDIS methodology to resting-state fMRI data with the simple modification of using anchor points at every possible volume within the dataset, i.e., from the first volume to the tenth from the last volume (whereafter, there would be insufficient volumes remaining for the spatial rearrangement of step 2, above) (**Figure 1B**). As in our previous work, to reduce computational costs, we first subsampled the input data to 4mm spatial resolution beforehand using the `fslmaths` command (– `subsamp2` option). The MELODIC command for step 5 was executed using the following settings: switch off brain extraction (BET), temporally-concatenated group-ICA; switch off MIGP (MELODIC’s Incremental Group-PCA) data reduction; dimensionality reduction set to 50 dimensions.

### Datasets and resting-state TARDIS execution

We downloaded cleaned MNI152 standard space resting-state fMRI data from the Human Connectome Project S1200 release (Feinberg et al., 2010). The scans were approximately 15 minutes each, acquired using a 3T Connectome scanner, with participants’ eyes open with relaxed fixation on a projected bright cross-hair on a dark background (and presented in a darkened room). The images were acquired with 2.0 mm isotropic voxels, TR=720ms (Chen et al., 2015; Feinberg et al., 2010; Setsompop et al., 2012). For our analyses we used the cleaned datasets for the first run, left-to-right (LR) encoding, of the first 50 participants (Van Essen et al., 2013).

All the TARDIS processing steps described above were executed on a high-performance cluster. With an input dataset of 50 resting-state fMRI scans (1200 volumes each), the final MELODIC command was executed on a compute node with 240gb RAM, taking ~48 hours to complete. When we present the TARDIS components, their numbering refers to the order (1..50) in which they were output from the MELODIC.

### Statistical analyses

To complement our qualitative assessment of the TARDIS components produced, we used a number of simple quantitative analyses. To examine the extent to which each TARDIS component was temporally variable within the component, we spatially correlated the fifth timepoint of each unthresholded component with all timepoints (1..10) from the same component using the `fslcc` command and an MNI152 T1 brain mask. The correlation values reported are therefore Pearson’s r coefficients. To provide a more quantitative and systematic interpretation of the TARDIS components, we used the Yeo cortical brain parcellation (Yeo et al., 2011) that defines seven ICNs. For each TARDIS component, we separately spatially correlated each of the seven Yeo ICNs with each timepoint (1..10) from within each unthresholded TARDIS component image using the `fslcc` command in a similar way. We recorded the maximum absolute r value between each ICN and every time point of each of the TARDIS components. For illustration purposes, a threshold of max(|r|) >0.4 was selected arbitrarily to denote high absolute spatial correlation.

To approximate how evenly each of the seven Yeo ICNs was maximally represented within each of the TARDIS components, for each component we calculated the Gini coefficient of the corresponding seven max(|r|) values. The Gini coefficient, ranging from 0 to 1, quantifies the degree of inequality in a distribution, from a situation of complete equality (=0, i.e. all Yeo components with the same max(|r|) value) to complete inequality (1, i.e. a single component with max(|r|) ~=0).

## Results

We computed 50 spatiotemporal TARDIS components from the cleaned 15-minute resting-state fMRI data of 50 HCP participants that had been registered into MNI152 standard space, using a TARDIS temporal window of ten fMRI volumes. These TARDIS components contain spatiotemporal patterns of signal covariance. A subset of the resulting components is shown in **Figure 2** and subsequently discussed; all 50 components are shown in **Supplementary Figure 1.**

**Figure 2.**
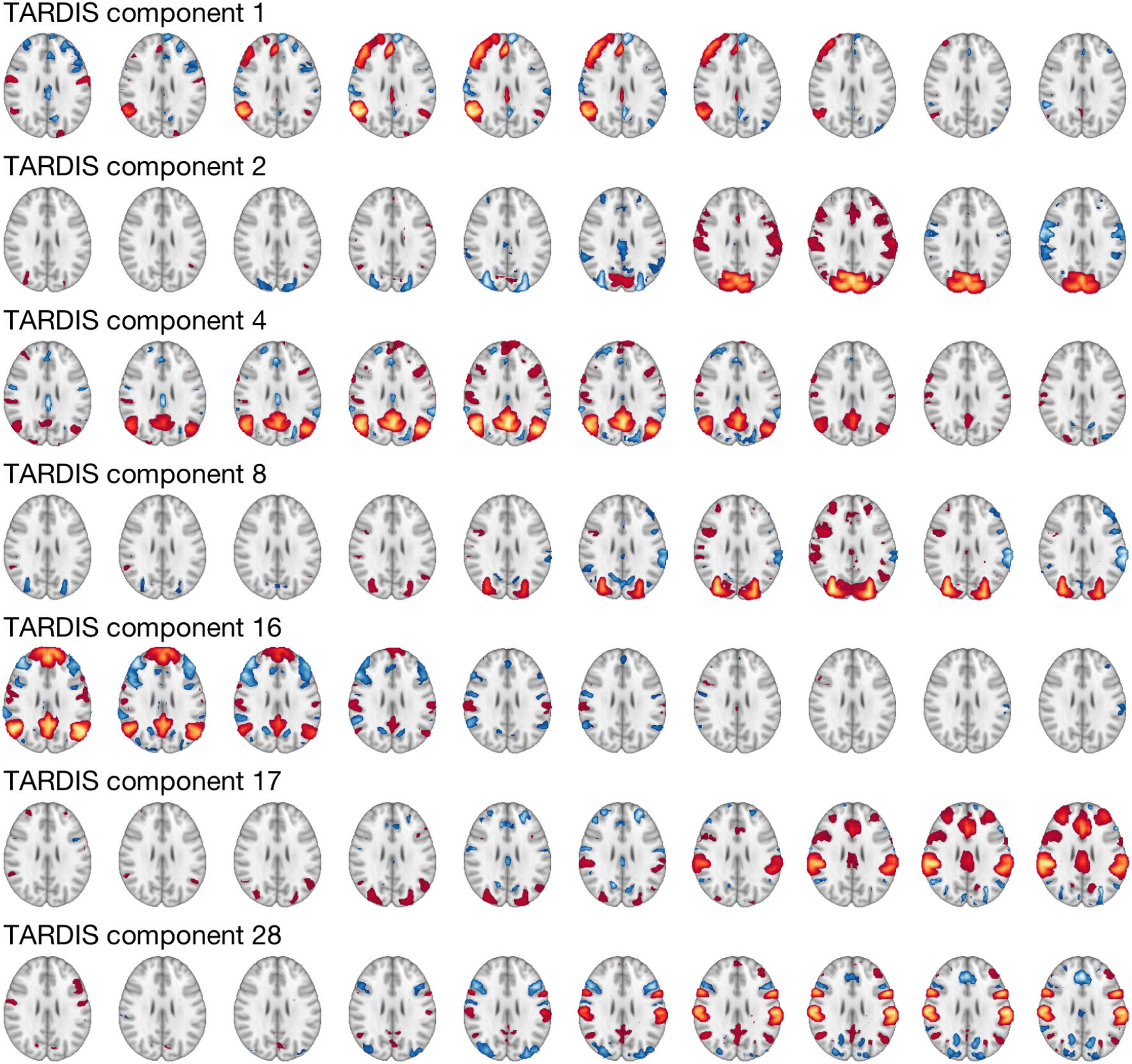
Selected spatiotemporal components identified using the TARDIS approach that illustrate the evolution over time of networks resembling canonical ICNs. Components are numbered according to their order (1..50) output from the MELODIC ICA (see Methods). Each component is shown as a row of axial slices, where columns are time (volume 1..10, left-right) within the component. Images are thresholded at an arbitrary |z|>2.3, with warm/cold colours indicating positive/negative weightings, respectively. The components are selected to illustrate examples of evolution of ICN-like networks over time within the component. See text for further description. All 50 components are shown in **Supplementary Figure 1**.

Qualitatively, many TARDIS components were spatially similar to canonical ICNs, spanning sensory and heteromodal brain regions. However, rather than a static snapshot, we observed changes in the spatial structure of components over time (i.e., from volumes one to ten). For example, we found within TARDIS component 1 the gradual onset and offset of a left-lateralized FPCN-like network. This temporal evolution is consistent with the typical cortical haemodynamic response function of approximately ~4-6 seconds (Taylor et al., 2018). In addition, we found that the onset of this lateralized FPCN-like network was preceded by a distributed pattern of both increases and decreases in other lateral and medial fronto-parietal regions.

We found several other TARDIS components also contained the onset and/or offset of other canonical ICN-like networks, such as components 2 and 8, both suggesting anti-correlated primary and lateral visual association networks, but each with different temporal evolutions. Components 4 and 16 were both consistent with a DMN, but with subtly different spatial distributions and time courses: component 4 showed the gradual onset and offset of this DMN-like network, accompanied by anti-correlated FPCN regions; component 16 also contained anti-correlated DMN and fronto-parietal regions, but with a distinct temporal evolution to component 4.

Also shown in **Figure 2**, in components 17 and 28 we observe the gradual onset of frontal and parietal regions, which are shared across both components. However, there are other regions which are inversely weighted between the components, e.g. the ventromedial prefrontal cortex (vmPFC) and some occipital regions. Equally, components 17 and 28 begin with different spatial patterns of activity. Contrasting these components demonstrates how our spatiotemporal decomposition can discover spatial patterns that are partially overlapping yet markedly different in how they covary with certain brain regions (e.g., the vmPFC), differing also in terms of their relationships to prior spatial patterns.

In addition to other networks with a similar spatial distribution to previously reported ICNs, we observed a number of putative non-neural noise components, shown and labelled in **Supplementary Figure 1**. The classification of these components as non-neural we based on either their spatial patterns (e.g., across white matter), or temporal progression (e.g. rapid changes that are neurobiologically implausible, given the known temporal properties of the neural BOLD signal).

We next examined the extent to which each TARDIS component was temporally variable by spatially correlating the fifth timepoint of each TARDIS component with all timepoints within the same component (**Figure 3**). We found a range of temporal profiles of correlation, with many components showing a profile of gradual change consistent with the expected dynamics of the BOLD signal, i.e., progression over seconds, with the largest changes over ~4-6 seconds (Taylor et al., 2018). In contrast, we observed a range of other temporal profiles for what are likely non-neural noise TARDIS components (labelled in **Figure 3** and, accordingly, in **Supplementary Figure 1**), varying from a stationary spatial distribution across the time window, relatively unstructured spatial distributions across the time window and even the possibility of a quasi-oscillatory signal across time (i.e., component 42, see **Supplementary Figure 1**).

**Figure 3.**
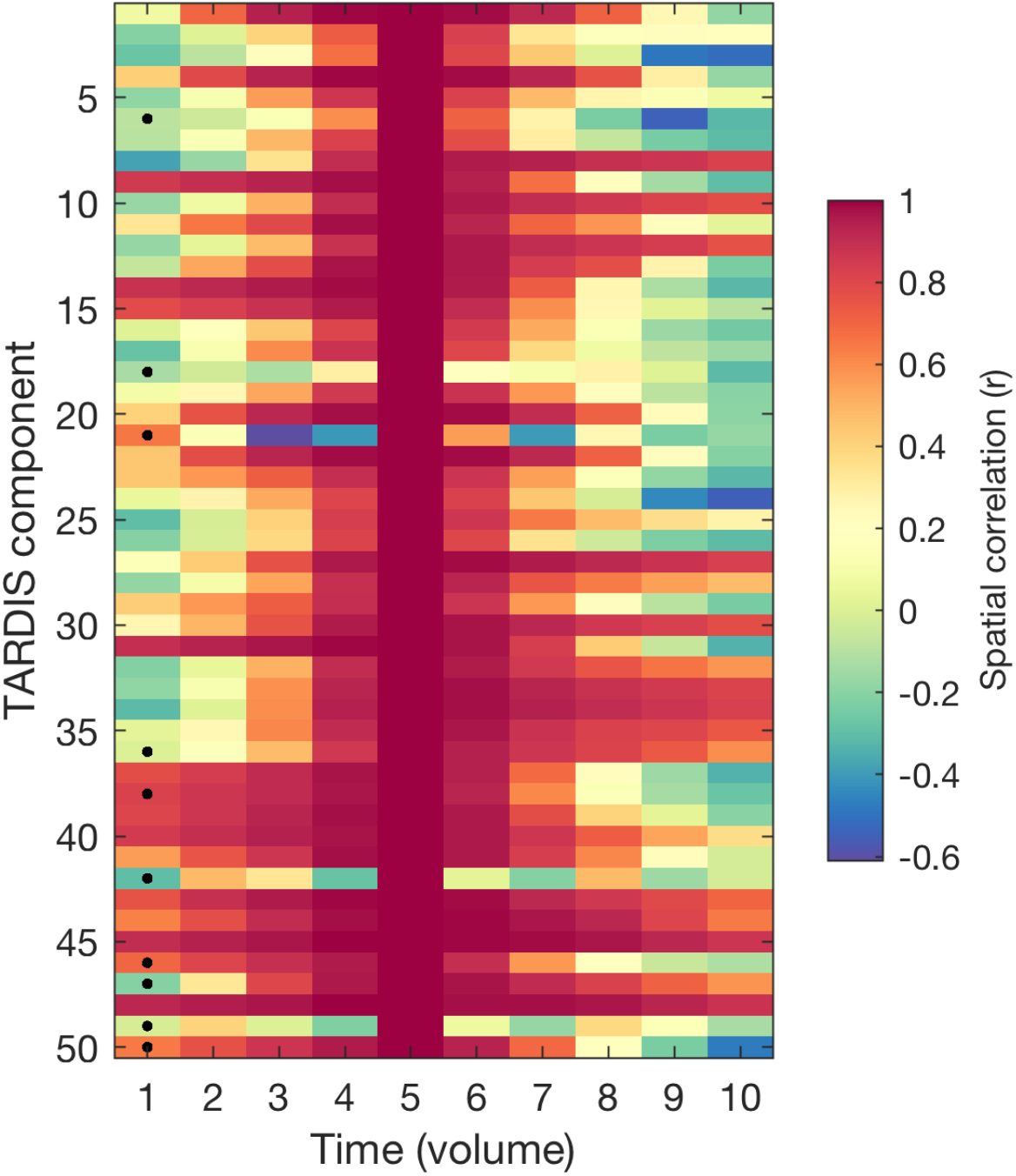
Temporal non-stationarity of TARDIS components. The spatial correlation coefficient (Pearson’s r) over time (left-to-right) of the nth timepoint (1..10) versus timepoint 5 is shown for each of the 50 components (see also **Figure 2**, see **Supplementary Figure 1** for all components). Components (rows) marked with a black dot indicate presumed non-neural noise components (see text). These components are labelled with an asterix (*) in **Supplementary Figure 1**.

To provide a more systematic and quantitative interpretation of the TARDIS components, we used the established Yeo cortical brain parcellation, based on resting state fMRI data (Yeo et al., 2011), and spatially correlated each timepoint of each TARDIS component with each of the seven canonical ICNs defined by Yeo et al. The results are presented in **Figure 4**, showing the seven Yeo ICNs (**Figure 4A**) alongside the maximum absolute value of spatial correlation (Pearson’s r) between each ICN and every timepoint of the 50 TARDIS components (**Figure 4B**). We found that each Yeo network, with the exception of the “limbic” network (terminology from Yeo et al), appears at some timepoint in one or more TARDIS components, i.e., with a high spatial correlation. (Bars in **Figure 4B** are shaded in colour wherever max(|r|)>0.4.) For example, we see that a spatial pattern similar (i.e., max(|r|)>0.4) to the Yeo-defined DMN occurs in four of the TARDIS components (**Figure 4B,** fourth row), with a maximum correlation coefficient within component 16 (see **Figure 2**),

**Figure 4.**
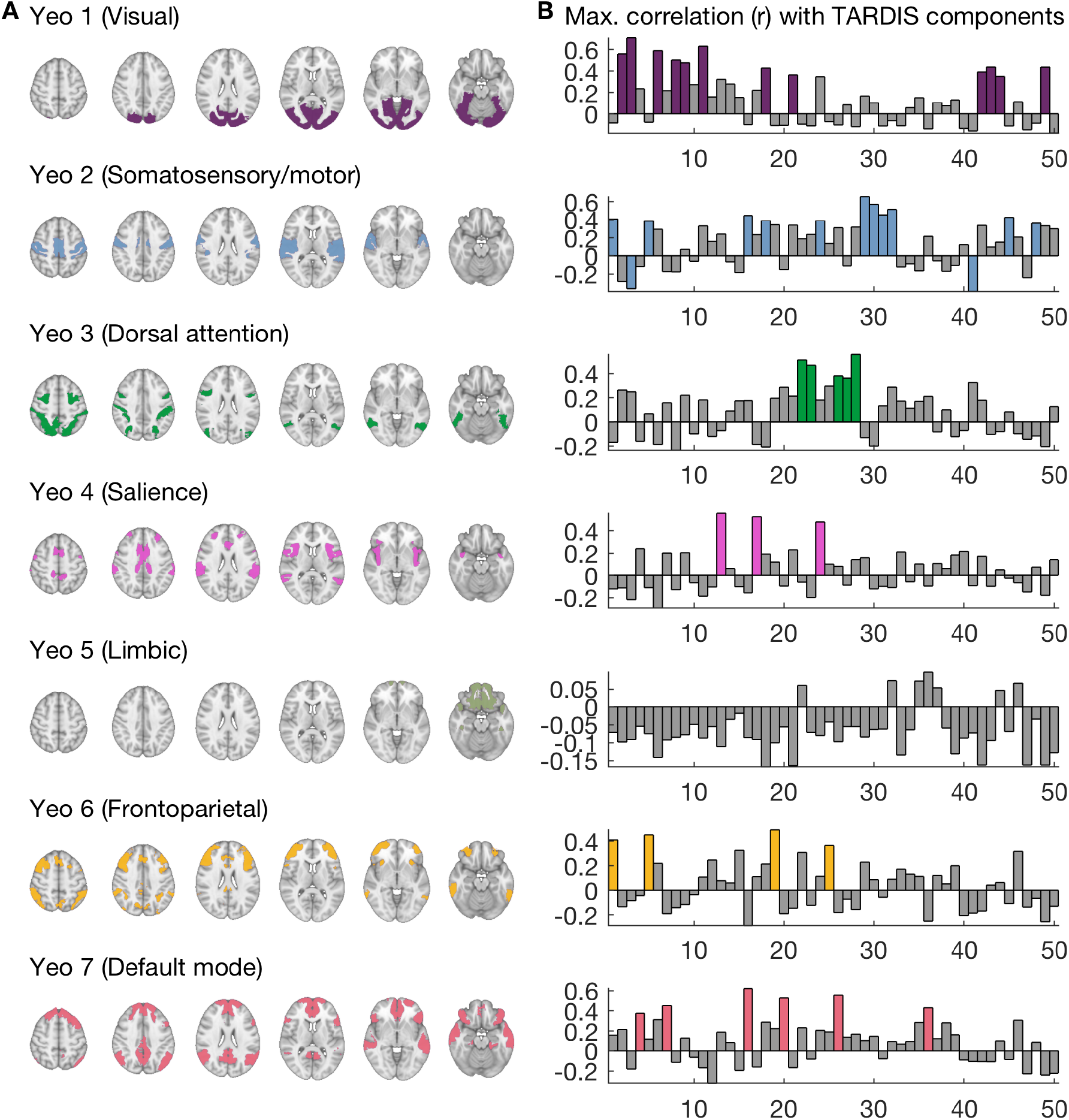
The set of seven Yeo reference ICNs and maximum correlation with TARDIS components. **(A)** The seven networks defined by Yeo et al using resting-state fMRI (Yeo et al., 2011). Colouring assigned to each network is arbitrary. **(B)** Maximum absolute spatial correlation (Pearson’s r, y-axis) between each timepoint (1..10) of each TARDIS component (1..50, x-axis), for each of the Yeo reference networks in (A). Coloured bars indicate components for which |r|>0.4 for one or more timepoints. Component numbering (x-axes) is the same as for all other figures.

We observed that most TARDIS components did not spatially correlate substantially with several Yeo networks, i.e. there was typically a predominance of one, or rarely two, ICNs, within components. The median Gini coefficient (0..1) for each non-noise TARDIS component, quantifying the equality of the maximum absolute correlation coefficients of each Yeo ICN within the component’s timepoints, was 0.41 (range 0.14-0.48). The Gini coefficients for all components are included in **Supplementary Figure 2.** For example TARDIS component 32, with the highest Gini coefficient, featured strong correlations (maximum r=0.52) only with the Somatosensory Yeo network (**Figure 4A**); in contrast, component 39, with the lowest coefficient, showed (weak) maximum correlations with all ICNs (0.11-0.2) (**Supplementary Figure 2**).

We finally used the Yeo parcellation ICNs (**Figure 4A**) as a framework to describe the temporal evolution of TARDIS components in further detail. Here we focus on a subset of components that relate to the DMN (Yeo network 7), presented in **Figure 5**, but all 50 components are shown using this decomposition in **Supplementary Figure 2**. The spatial map at each timepoint of each TARDIS component had been correlated with each of the seven Yeo networks. Using this method of description, we observed a range of DMN-related spatiotemporal dynamics. For example, in TARDIS components 4 and 7 (**Figure 5A** and **B**), we observed the onset of a DMN-like network, i.e., increasing spatial correlation with Yeo network 7, but with different times of onset (i.e. occurring later in component 7, from volume ~6 to 10) and involving different anti-correlated regions. In contrast to these DMN onsets, in component 16 (**Figure 5C**) we observe the offset of the DMN from the beginning of the component, followed by the onset of Yeo network 2 (somatosensory/motor). In component 20 (**Figure 5D**), the DMN onset and subsequent offset is flanked by the onset of Yeo network 3 (dorsal attention). In components 26 and 36 (**5D&E**), we again found onsets of DMN-like networks, but these were associated with differing evolution of other networks.

**Figure 5.**
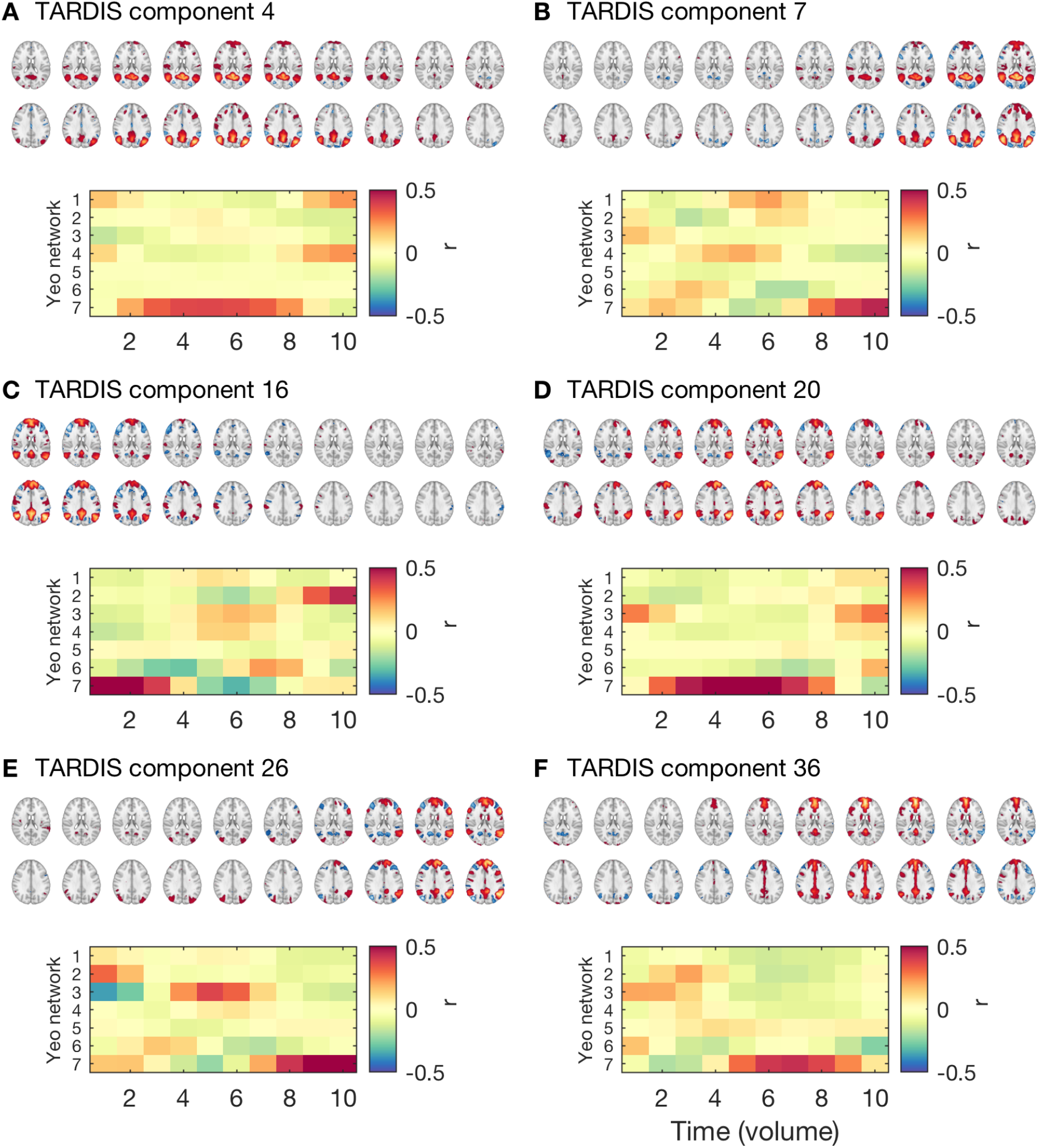
Spatiotemporal dynamics of DMN-like patterns within TARDIS components. Each TARDIS component is shown as two rows of axial slices, where columns are time (volume 1..10) within the component. Dynamics are quantified in terms of spatial correlations (Pearson’s r between each of the seven Yeo ICNs (Yeo et al., 2011) (**Figure 4A**) at each timepoint (1..10) of each TARDIS component separately (seven rows by ten columns). Panels **(A-F)** Show selected components each involving a DMN-like network, i.e. containing a timepoint with high spatial correlation with Yeo network 7 (**Figure 4A**). Component numbering is the same as in **Figures 2–4**. Colour bars are clamped to +/− 0.5.

In general, we observe that the spatiotemporal decomposition of the TARDIS components was consistent with the spatial distribution captured by the Yeo networks; individual timepoints of the TARDIS components were each correlated with a unique Yeo network, rather than being a blend between multiple Yeo networks, or appearing as a distinct spatial pattern.

## Discussion

We have applied a data-driven approach, sensitive to both spatial and temporal non-stationarity, to resting-state fMRI in order to discover spatiotemporal patterns. We observed that the components recovered could generally be described in terms of intrinsic connectivity networks (ICNs) that have been previously calculated based on techniques that assumed spatial stationarity: our spatiotemporal components could be characterised as graded onsets and offsets of these well-described networks. Our results suggest that these ICNs are relatively spatially stable when they occur in resting-state fMRI, whether they are increasing or decreasing in activity and whether such changes are following or preceding other networks. This implies that, to a reasonable approximation, stable ICNs can be taken to be building blocks of the spatiotemporal patterns measured with resting-state fMRI.

Our TARDIS approach involves a simple modification of the input data, subsequently passed through a spatial ICA used frequently in resting-state fMRI analysis (i.e., FSL MELODIC). As such, the assumptions and benefits of our approach are similar to the classic data-decomposition techniques that are very familiar in functional imaging. The only additional constraint is the temporal window for the spatial remapping, which limits for how long spatiotemporal dynamics (and potential non-stationarity) can be observed. We set this window to be longer than the haemodynamic response (~4-6s) typically reported in fMRI.

One important difference with standard spatial ICA applied to fMRI is that different components capture covariance in signals at specific spatial (i.e., voxel) *and* temporal positions; so two patterns that could be spatially similar enough to be grouped together into a single component with typical spatial ICA, will be in different components depending on when they happen. That is, for two putative neural patterns A and B, A following B will be in a separate component from B following A. Our simple approach, therefore, allows us to investigate both the hypothesis where A following B is matched by B following A (spatial stationarity), from the non-stationary hypothesis e.g., B is followed by a slightly different pattern A’. The results from our analysis suggest the former, stationary, case predominates.

Most of the neural components we observed had spatial patterns with graded onsets and offsets of networks that spatially correlated with the Yeo networks. This is the type of pattern that might be expected if resting-state fMRI involves transitions between a relatively small number of stable networks. This is a reassuring finding for the large number of studies that have assumed spatial stationarity and reported transitions between neural states as important for understanding a range of aspects of resting-state fMRI data, and that have related these states to other factors such as ongoing thought reported from experience sampling, or clinical conditions, e.g., (Karapanagiotidis et al., 2020).

We did not observe components clearly indicating spatial non-stationarity over time. For instance, we did not observe fractionated versions of the ICN-like networks, beyond simple left-right homology, within components. Neither did we observe spatial spread of an ICN-like network over time to involve regions outside the network. This does not preclude the existence of more subtle spatial non-stationarities; there could be situations that are less common, or less consistent, or of much smaller magnitude, such that they do not feature prominently in our ICA-derived components. Alternative approaches, such as using recurrent or transformer neural networks may be able to pull out clear examples of spatial non-stationarity. That said, our results do provide evidence that, at least to a first approximation, resting-state fMRI data can be relatively well described as a sequence of transitions between, or global modulations of, spatially static canonical ICNs.

Although we have focused on the components that, given their spatial and temporal properties, are more likely neural, the components we designated as non-neural noise are also potentially of interest. These spatiotemporal noise components may better capture and identify non-neural variance in the data compared to traditional methods. One avenue for future work is to investigate whether filtering out these components could clean fMRI data and improve subsequent analyses, even those subsequently using traditional analysis, e.g., seed-based approaches.

There are a number of limitations arising from the current work. First, the use of data transformation (remapping time into space) before ICA means that our results are most comparable to other approaches using spatial ICA; other techniques for data reduction (e.g., principle component analysis (PCA), diffusion embedding, etc.) are likely to result in somewhat different spatiotemporal patterns. In addition, the ICA approach is probabilistic, and a different random seed would result in subtly different spatiotemporal patterns; we note, however, that in our work, we explored this variability across different dimensionalities of data reduction and found broadly consistent results. The approach operates across all voxels, as such, there may be more subtle patterns of spatiotemporal non-stationarity that are missed at the whole-brain level and may only be apparent at the regional or voxel scale or with the higher granularity from extracting more components. Finally, the expanding of the ICA input data (by close to a factor of 10) places a substantial computational burden, increasing processing time and limiting the length of e.g., the temporal window considered.

Many commonly used fMRI analysis techniques, such as whole timeseries temporal correlation between regions or voxels, impose spatial stationarity; they also impose the requirement of non-overlapping results (i.e., each voxel has a single correlation with every other voxel). Approaches based on sliding windows or more sophisticated extensions, e.g., (Monti et al., 2014), reduce the requirement of stationarity to the length of the window. Alternative data decomposition approaches (e.g., spatial ICA, PCA, diffusion embedding, etc.) do allow for the discovery of graded responses, with voxels having multiple relationships with other voxels; however, they typically apply to the whole time series or windowed time series. Our TARDIS approach, within the constraints of the number of timepoints considered (ten in this case), allows for both spatial and temporal non-stationarity, as well as the ability to find multiple potentially spatially and temporally overlapping components. In this initial application to resting-state fMRI data, we were able to derive spatiotemporal components in a data-driven way, and examine how these components related to a commonly used framework for describing resting-state fMRI, the parcellation of Yeo et al. This approach suggests a way to explicitly test how well these frameworks work in capturing the variability in resting-state fMRI data, and how much of a role more complicated, non-temporally and non-spatially stationary patterns of brain activity play in macroscopic neural activity.

## Conclusion

Our analyses suggest that the spatiotemporal evolution seen within TARDIS components primarily resemble transitions between previously-described canonical brain networks (ICNs). However, this does not rule out the possibility that there are spatial patterns which account for a smaller proportion of the variance in the data, that do not correspond to canonical networks, for example, relatively subtle voxelwise differences between two components that are nonetheless somewhat similar at a coarse level. This suggests that a relatively consistent, spatially stationary, set of core networks underlies the majority of the spatiotemporal dynamics we observed.

## Acknowledgements

GS is funded by the National Institute for Health Research (NIHR). Data were provided [in part] by the Human Connectome Project, WU-Minn Consortium (Principal Investigators: David Van Essen and Kamil Ugurbil; 1U54MH091657) funded by the 16 NIH Institutes and Centers that support the NIH Blueprint for Neuroscience Research; and by the McDonnell Center for Systems Neuroscience at Washington University.

## Author contributions

Both authors were responsible for conception and design of the study and writing the manuscript. GS was responsible for carrying out the analysis of data.

## Supplementary Materials

The two supplementary figures appear on the following two enlarged pages.

**Supplementary Figure 1.**
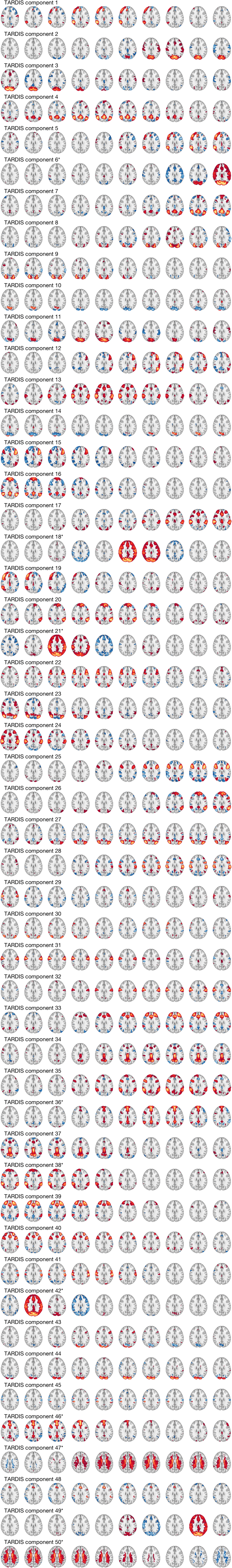
The 50 spatiotemporal components identified using the TARDIS approach. Components are numbered according to their order (1..50) output from the MELODIC ICA (see Methods). Each component is shown as a row of axial slices, where columns are time (volume 1..10, left-right) within the component. Images are thresholded at an arbitrary |z|>2.3, with warm/cold colours indicating positive/negative weightings, respectively. Component numbers labelled with an asterix (*) indicate putative non-neural noise components, based on their distributed spatial pattern (e.g., across white matter), or temporal progression (e.g. rapid changes that are neurobiologically implausible, given the known temporal properties of the BOLD signal).

**Supplementary Figure 2.**
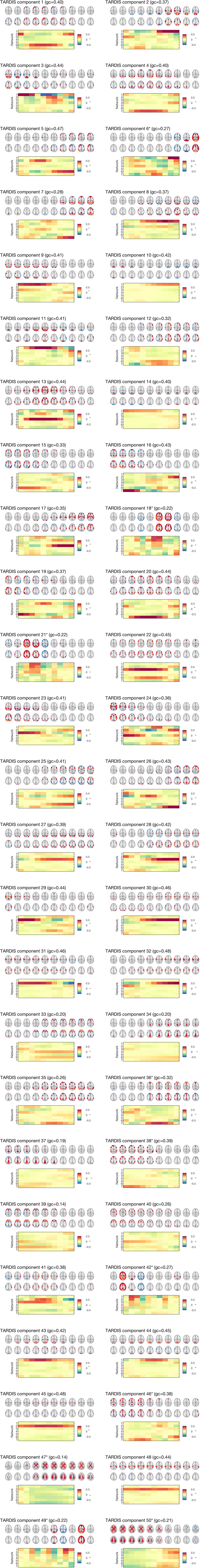
Spatiotemporal dynamics within TARDIS components. Each TARDIS component is shown as two rows of axial slices, where columns are time (volume 1..10) within the component. Dynamics are quantified in terms of spatial correlations between each of the seven Yeo ICNs at each timepoint (1..10) of each TARDIS component separately (seven rows by ten columns). For TARDIS each component, the Gini coefficient (gc) of the maximum spatial correlation (max(|r|) of each Yeo ICN across the timepoints of the component, is shown.

